# VIGA: a sensitive, precise and automatic *de novo* VIral Genome Annotator

**DOI:** 10.1101/277509

**Authors:** Enrique González-Tortuero, Thomas David Sean Sutton, Vimalkumar Velayudhan, Andrey Nikolaevich Shkoporov, Lorraine Anne Draper, Stephen Robert Stockdale, Reynolds Paul Ross, Colin Hill

## Abstract

Viral (meta)genomics is a rapidly growing field of study that is hampered by an inability to annotate the majority of viral sequences; therefore, the development of new bioinformatic approaches is very important. Here, we present a new automatic *de novo* genome annotation pipeline, called VIGA, to annotate prokaryotic and eukaryotic viral sequences from (meta)genomic studies. VIGA was benchmarked on a database of known viral genomes and a viral metagenomics case study. VIGA generated the most accurate outputs according to the number of coding sequences and their coordinates, outputs also had a lower number of non-informative annotations compared to other programs.

## Introduction

Virology is a diverse scientific discipline. While many researchers are interested in discovering and characterising pathogenic eukaryotic viruses [1], recently there has been an increased interest in revealing bacteria- and archaea-infecting viral communities [2]. The number of viral metagenomic studies is increasing due to the development of new sequencing technologies and the reduction in costs. However, due to the volume of information that these platforms generate and the large proportion of viral sequences sharing little or no homology to known viral genomes (‘viral dark matter’, [3]), new bioinformatic tools are required to examine viral contigs and genomes [4].

Viral annotation methods differ depending on the host organism. Bacteriophages and archaeal viruses are annotated using prokaryotic genome annotation software or web-servers such as RAST [5], Prokka [6] and RASTtk [7]. However, these bioinformatic tools are optimised for bacterial sequences, not viruses (despite the improvements in RASTtk to annotate phage sequences [8]). In contrast, eukaryotic viruses are annotated using close-reference based methods such as FLAN [9], VIGOR [10] and ViPR [11]. In a similar way, VirSorter [12] and VirusSeeker [13] were designed to predict putative prokaryotic viral contigs in metagenomic datasets. However, both programs predict viral contigs according to the presence of viral proteins using reference databases, and close-reference homology-based methods can underestimate true viral diversity due to database limitations [3,14]. Therefore, in this manuscript, we present a new modular and automatic *de novo* genome annotation bioinformatic pipeline, called VIGA (VIral Genome Annotator), to annotate viral sequences.

VIGA automatically detects open reading frames from a FASTA or multi-FASTA formatted file. VIGA then annotates protein sequences by detecting homologues in a BLAST (“Slow”) or a DIAMOND (“Fast”) protein database, with or without Hidden Markov Model (HMM) protein detection against a protein database. The different methodologies for annotating viral contigs and genomes allows the user to specify options that sacrifice annotation detail in exchange for increased speed, which is required when dealing with larger metagenomic datasets. In addition, VIGA also automatically detects (1) the topology of viral contigs, (2) the presence of rRNA, tRNA and tmRNA sequences, (3) potential CRISPR repeats and (4) tandem or inverted repeat sequences. Finally, VIGA outputs a FASTA file that includes user specified modifiers, a GenBank file and a five-column tab-delimited feature file to ease the upload of annotated contigs and genomes to various database repositories and genome visualisation platforms.

## Results

### Benchmarking of VIGA

The performance of VIGA, Prokka, RAST and RASTtk was tested using a benchmark database comprising 191 sequences belonging to 138 different viruses (52 bacteriophages, 72 eukaryotic and 10 archaeal viruses, and 4 virophages; Additional file 1). Of the 72 eukaryotic viruses, 11 have multipartite genomes. Experimental evidence is available for the coding sequences of 117 out of the 123 sequences of eukaryotic viruses, 28 out of 52 sequences of bacteriophages, 3 out of 10 sequences of archaeal viruses, and none of the 4 virophage sequences used. When bioinformatic methods were used to annotate these viral genomes in the original data, a wide variety of methods were employed, including GeneMark [15], GLIMMER [16], NCBI ORF Finder and the University of Wisconsin Genetics Computer Group [17]. The outputs of VIGA, Prokka, RAST and RASTtk were evaluated according to three different parameters: (1) number of coding sequences, (2) coordinates of coding sequences, and (3) power of prediction.

Firstly, the accuracy and the precision of the number of viral coding sequences were estimated using general linear models. Accuracy was measured by the slope, and precision was measured according to the coefficient of determination (R^2^). To compare all these linear models, analysis of covariance (ANCOVA) was performed. In a general overview, the programs delivered different estimates of the number of coding sequences (ANCOVA: *p* < 2×10^−16^). In fact, although all programs tended to overestimate the number of genes, VIGA provided the most accurate predictions (i.e. accuracy is closest to one, Fig. 1A). Moreover, VIGA and Prokka had very similar values of precision (Table 1). When compared according to viral host, similar results were found only in the case of eukaryotic viruses (ANCOVA (Archaeal viruses): *p* = 0.922; ANCOVA (Bacteriophages): *p* = 0.734; ANCOVA (Eukaryotic viruses): *p* = 1.560×10^−15^; Figs. 1B-D). Interestingly, when bacteriophages were considered, only RASTtk tended to overestimate the number of coding sequences (Table 1).

**Table 1.** Accuracy and precision in the number of coding sequences

| Case | Program | Accuracy<br>(Slope) | Precision<br>(R<sup>2</sup>) |
| --- | --- | --- | --- |
| **General** | *VIGA* | 1.027 | 0.997 |
|  | *Prokka* | 1.043 | 0.996 |
|  | *RAST* | 1.118 | 0.979 |
|  | *RASTtk* | 1.135 | 0.982 |
| **Archaeal viruses** | *VIGA* | 0.962 | 0.990 |
|  | *Prokka* | 0.991 | 0.991 |
|  | *RAST* | 0.821 | 0.936 |
|  | *RASTtk* | 1.036 | 0.993 |
| **Bacteriophages** | *VIGA* | 0.997 | 0.997 |
|  | *Prokka* | 0.983 | 0.995 |
|  | *RAST* | 0.982 | 0.996 |
|  | *RASTtk* | 1.015 | 0.997 |
| **Eukaryotic viruses** | *VIGA* | 1.031 | 0.997 |
|  | *Prokka* | 1.050 | 0.997 |
|  | *RAST* | 1.136 | 0.979 |
|  | *RASTtk* | 1.151 | 0.982 |

**Figure 1.**
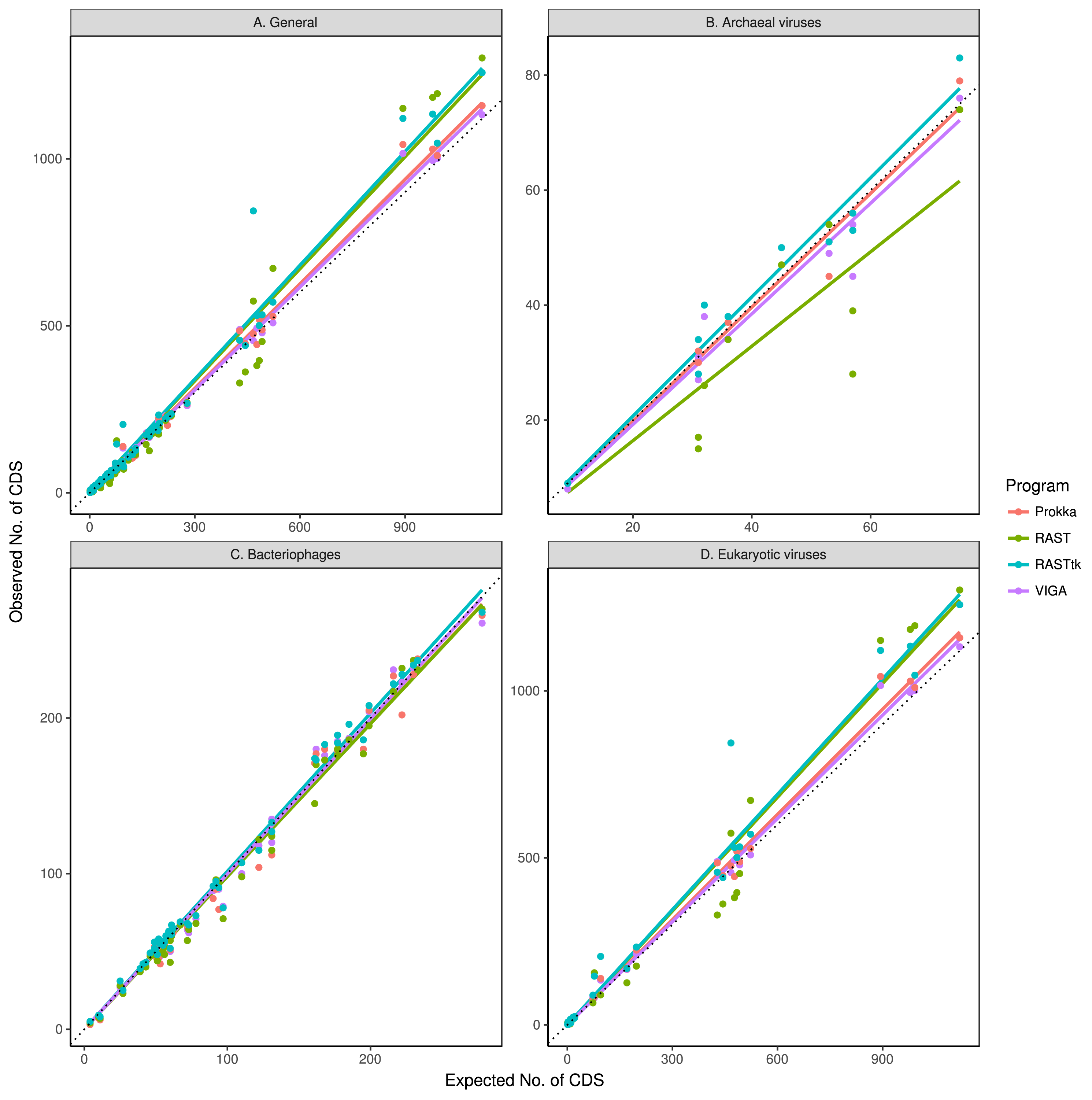
Correlation between the expected and observed number of coding sequences when considering (A) all known viral sequences, (B) archaeal viruses, (C) bacteriophages, and (D) eukaryotic viruses. Dotted line is a 1:1 line.

Secondly, F_1_ score, a measure that combines precision and sensitivity, was used to predict the quality of the coordinates of the viral coding sequences. Moreover, to evaluate the occurrence of false positives (i.e. false coordinates considered as true; type I error) and false negatives (i.e. true coordinates considered as false; type II error), false discovery rate (FDR) and false negative rate (FNR) were examined. VIGA scored very highly for both bacteriophages and eukaryotic viruses. In eukaryotic viruses the highest false discovery rate (FDR) was associated with RASTtk, while RAST had the highest false negative rate (FNR). For bacteriophages the highest FDR and FNR were obtained for Prokka. In the case of archaeal viruses, VIGA again had the highest precision, while the highest sensitivity was noted in RASTtk (Table 2).

**Table 2.**
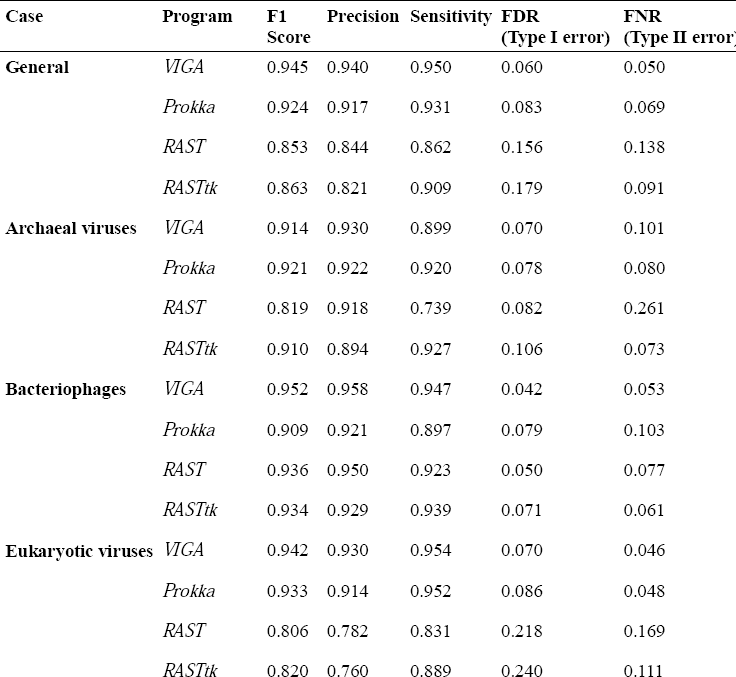
Accuracy, precision and sensitivity of the different programs. False Discovery Rate (FDR) and False Negative Ratio (FNR) are used to describe errors in the precision and sensitivity.

Finally, the power of prediction of all programs was measured by considering the number of non-informative annotations (i.e. all proteins classified as “hypothetical protein”, “uncharacterized protein”, “ORF”, “predicted protein”, “unnamed product protein” or “gp[Number]”). For these analyses, two different modes of running VIGA were considered – “Slow” (when BLAST and HMMER are used to annotate the genes) and “Fast” (when DIAMOND alone is used for annotation). Kruskal-Wallis (KW) test was performed to detect potential differences in the power of prediction of all three programs (including both variants of VIGA) and significant differences between the various programs were observed (KW test: *p* = 1.683×10^−53^). In all cases, no significant differences between VIGA-Slow and VIGA-Fast were found (Nemenyi test: *p* = 0.853). In fact, while RAST and RASTtk had the highest number of non-informative annotations, both VIGA modes had the smallest number (Fig. 2A). Additionally, there were significant differences among programs independently of the viral type (Table 3). In all cases, VIGA achieved optimal annotation, having always the smallest number of non-informative annotations. In contrast, Prokka had the highest amount of non-informative annotations in prokaryotic viruses (Figs. 2B-C) and RAST and RASTtk had the highest amount of non-informative descriptions in eukaryotic viruses (Fig. 2D).

**Table 3.**
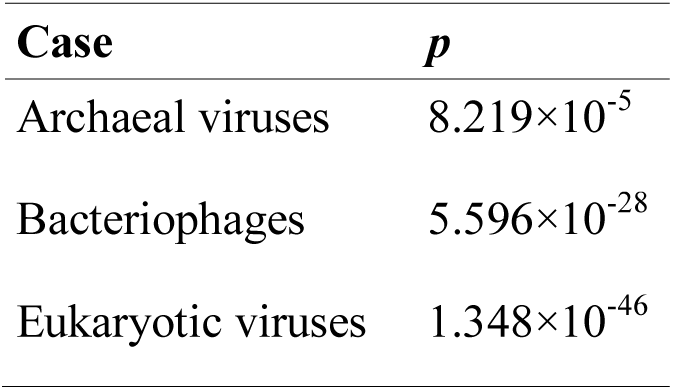
Kruskal-Wallis *p*-values for the comparison between all different pipelines considering the different viral types.

| Case | $p$ |
| --- | --- |
| Archaeal viruses | $8.219 \times 10^{-5}$ |
| Bacteriophages | $5.596 \times 10^{-28}$ |
| Eukaryotic viruses | $1.348 \times 10^{-46}$ |

**Figure 2.**
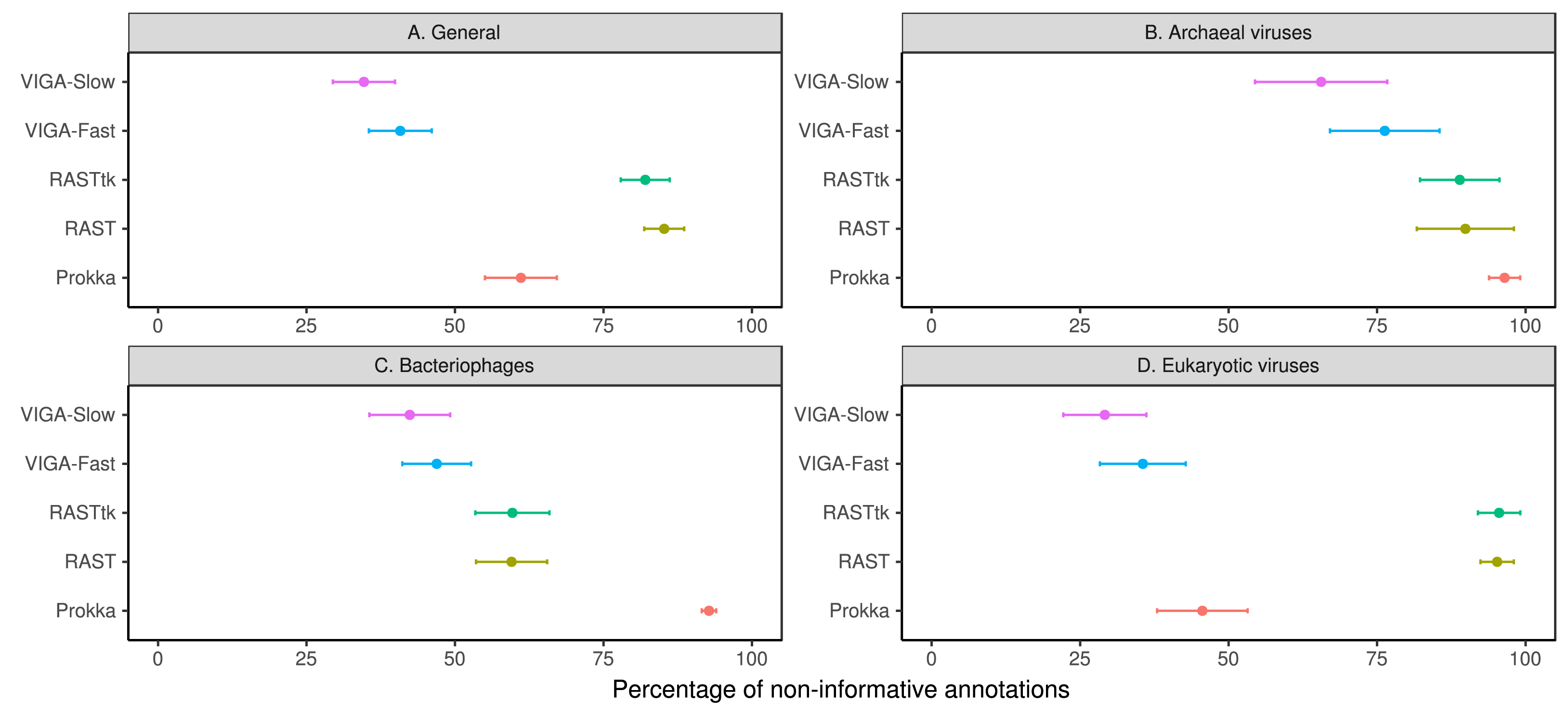
Percentage of non-informative annotations when processed in all programs for (A) all known viral sequences, (B) archaeal viruses, (C) bacteriophages, and (D) eukaryotic viruses. Dot indicates the average value of non-informative annotations and bars indicates the 95% confidence interval.

### Case study: healthy human gut phageome

To evaluate the performance of VIGA on a metagenomic dataset, VIGA, Prokka, RAST and RASTtk were run using a subset of 202 non-redundant contigs from the metavirome of healthy individuals [18]. VIGA was executed using 10 cores in two different ways: (1) using only DIAMOND (VIGA-Fast), and (2) using BLAST and HMMER (VIGA-Slow). These 202 contigs were composed of 65 short contigs (<15 kb), 99 medium-size contigs (15 – 70 kb), and 38 long contigs (>70 kb). Two different parameters were evaluated: (1) Speed of the program, and (2) power of prediction. Only RASTtk was unable to annotate these contigs.

To test the speed of VIGA-Slow and VIGA-Fast, both VIGA modes and Prokka were run in a local server (Lenovo x3650 M5, with 48 Intel Xeon 2.6GHz Processors, Ubuntu 14.04, 512 GB of RAM) using 10 processors. VIGA-Slow took 19,283 minutes (13 days 9 hours 23 minutes) to process all 202 contigs of this data set, while VIGA-Fast took 809 minutes (13 hours 29 minutes). In contrast, Prokka took 3 minutes to annotate all contigs. Unfortunately, we cannot estimate the time that RAST took to annotate these genomes due to be an external web server.

Finally, the power of prediction of all programs was evaluated by comparing the number of non-informative annotations as indicated above. Significant differences between the various programs were observed (KW test: *p* = 2.121×10^−93^). While Prokka had the highest percentage of non-informative descriptions, VIGA-Slow had the smallest number (Fig. 3A). In contrast to the benchmark, there were significant differences between VIGA-Slow and VIGA-Fast on a metagenomic dataset. VIGA-FAST had a higher percentage of non-informative descriptions than VIGA-Slow (Nemenyi test: *p* = 3.900×10^−14^). Surprisingly, no significant differences between VIGA-Fast and RAST were found (Nemenyi test: *p* = 0.440; Fig. 3A). When the different size of contigs were considered, significant differences between the non-informative annotations of the programs were found (KW test (“Short”): *p* = 4.650×10^−24^; KW test (“Medium”): *p* = 3.731×10^−63^; KW test (“Long”): *p* = 8.708×10^−16^). This is a similar pattern detected independently of the contig size (Figs. 3B-D).

**Figure 3.**
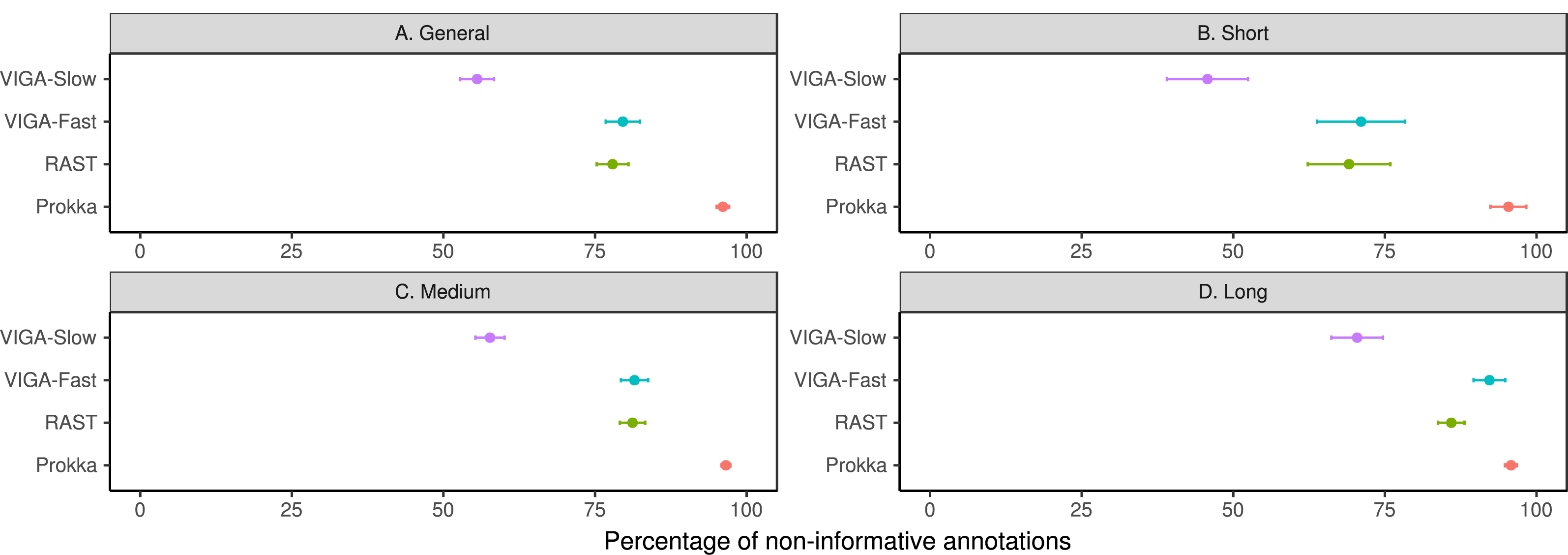
Percentage of non-informative annotations for the case study dataset when processed in all programs for (A) the case study dataset, (B) short contigs (<15 kb), (C) medium contigs (15 – 70 kb), and (D) long contigs (>70 kb). Dot indicates the average value of non-informative annotations and bars indicates the 95% confidence interval.

## Discussion

In this study, VIGA, a new bioinformatic pipeline for viral genome annotation, was tested against RAST, RASTtk and Prokka using a benchmark comprising of 138 viruses. In fact, this is the first genome annotation pipeline to be benchmarked using viral data, as previous validation of these programs tended towards the use of bacterial genomes [5,6]. When all these bioinformatic annotation pipelines were benchmarked, VIGA successfully outperformed the others in all test parameters. After validating VIGA, it was used to annotate the phages in a subset of the Manrique et al. healthy human gut phageome dataset [18]. This subset was based on the phages predicted by VirSorter [12], which could miss some viral contigs such as variants of crAssphage [19]. In that instance, this viral gene annotation is dependant on the proficiency of VirSorter.

When the benchmark of 138 viruses was performed to measure the accuracy and precision of the number of coding sequences, VIGA had the highest values of accuracy and precision in the general overview. The only differences in the number of coding sequences were shown in eukaryotic viruses. Additionally, when the quality of the coordinates of these coding sequences was analysed, RASTtk had the highest false discovery rate and RAST the highest false negative rate for eukaryotic viruses. All these observations strengthen the idea that all tested programs were developed for prokaryotic viruses. Although the most abundant viruses in the biosphere are bacteriophages [20], it was not possible to annotate around 80% of putative viral contigs in previous studies on viral diversity [14], indicating the extensive presence of ‘viral dark matter’. The nature of this ‘viral dark matter’ is related with the lack of knowledge in viral diversity, and due to the use of homology-search methods to classify and to annotate them [3]. In that sense, classification of viruses (independently of their hosts) currently should not only be performed using close-reference based homology searches because they could underestimate the real viral diversity based on the limitations of databases.

The quality of the coordinates of the coding sequences in the viral benchmark was higher using VIGA than with the other programs. Although this result suggests that VIGA is reliable, it is also important to note that there was only experimental evidence of the coding sequences in 68 of 74 sequences of eukaryotic viruses, 28 of 52 sequences of bacteriophages, and 3 of 10 sequences of archaeal viruses. In fact, although the development of automatic genomic pipelines such as RASTtk or VIGA can facilitate the prediction of genes in viral sequences, some features such as introns, morons or regulatory elements need manual refinement [8]. For this reason, all bioinformatic genome annotations are putative until validated using experimental procedures such as cDNA-gDNA hybridization [21–23], proteomics [24–26] or transcriptomics [27–29].

Analysis of the power of prediction of annotation pipelines showed that RAST and RASTtk tend to generate a higher number of non-informative annotations, while VIGA had the smallest number in all cases. Therefore, VIGA-Slow mode has the potential to provide more information on encoded viral genes than other genome annotation bioinformatic pipelines, which rely exclusively on homology-based methods such as BLAST, BLAT [30] or DIAMOND. Primarily because these methods increase the number of non-informative annotations, especially in novel viruses, as demonstrated in the described metagenomic case study. Viral dark matter [3], or the unknown fraction of the virome, is a prevalent hurdle in virome research and lack of homology to sequences in databases hampers most annotation methods. It is also important to note, that where annotations are available, many have been generated through bioinformatics and do not have supporting experimental evidence. It is therefore very important to consider the source of functional information for proteins when annotating new viruses unless empirical evidence is available [8,31].

Proteins related to viral function can have highly conserved sequences, such as the hepatitis B virus core protein [32], Dengue virus polyprotein [33] and the influenza A virus nucleoprotein [34], because non-synonymous mutations in these proteins could hamper viral function. For this reason, the use of HMMs was implemented to predict the putative function of these genes. Use of HHPred or InterProScan is suggested to increase the power of protein annotation predictions [31,35,36]. Although the implementation of these programs could be beneficial for VIGA and it will be implemented in future versions, HMM-based methods are slower than homology searches as noted in the case study. Another alternative to these HMM-based methods could be the implementation of homology-independent annotation methods such as iVIREONS [37] or VIRALpro [38]. All these methods use machine learning to predict structural phage proteins such as capsid, collar and tail proteins [8] and are also scheduled for implementation in future versions of VIGA. Finally, when the power of prediction of all genome annotation pipelines was analysed, a lack of criteria for gene annotations was found, making it difficult to compare between the outputs of the different programs. For this reason, the implementation of a standardised genome annotation system would ease the comparison between genomes [39,40] using some (alpha)numerical classifications such as the Enzyme Codes [41], Clusters of Orthologous Groups [42], KEGG Orthology [43] or the Prokaryotic Viral Orthologous Groups [44] which could be added in the genome annotation output.

## Conclusions

The number of viral metagenomic studies is increasing as a consequence of the development of high throughput sequencing platforms and cost reductions. However, there are few software programs to annotate the viral sequences and never before have these programs been benchmarked against each other. In this study, we present VIGA, a new automatic *de novo* genome annotation bioinformatic pipeline to annotate prokaryotic and eukaryotic viral sequences from genomic and metagenomic studies. VIGA allows the most accurate, precise and sensitive annotation of viral genomes when benchmarked using 138 known viral genomes. VIGA can be executed using BLAST or DIAMOND to annotate proteins according to homology, with the option to also use HMMER to improve these annotations based on HMMs. The use of HMMs will enrich the annotation detail of the viral contigs, but will decrease the speed of the program. Where increased speed is required for example when dealing with larger metagenomics datasets.

## Materials and methods

### Workflow of the software

*Overview.* VIGA is an automatic *de novo* viral genome annotator implemented in Python 2.7 (requiring Biopython [45]) and designed to annotate complete and draft viral and phage genomes comprising single or multiple contigs (Fig. 4). As an input, VIGA accepts a DNA FASTA file with the (putative) viral contigs. These sequences are processed to predict the topology of the contigs (i.e. circular or linear). If the contig is circular, the prediction of the origin of replication is performed according to cumulative GC skew and realignment of the contig from the putative origin of replication. Coding sequences (CDS) are predicted and, then, the function of these proteins is inferred based on homology using BLAST [46] or DIAMOND [47] and, optionally, using Hidden Markov Models (HMMER [48]). After that, a decision tree algorithm chooses the most reliable description of the protein (Fig. 5). Potential rRNA sequences are predicted using INFERNAL [49] with the use of the Rfam database [50], and tRNA and tmRNA sequences are predicted using ARAGORN [51]. Additionally, CRISPR, tandem and inverted repeats are predicted using PILER-CR [52], Tandem Repeats Finder [53] and Inverted Repeats Finder [54] respectively. Repeat sequences are related with the gene expression regulation, integration of the viral genome and, even, viral replication. Finally, the output of the program are a GenBank file, a FASTA file and a table (TBL) file suitable for GenBank submission (Fig. 4). Optionally, a General Feature Format (GFF) version 3 file can be generated.

**Figure 4.**
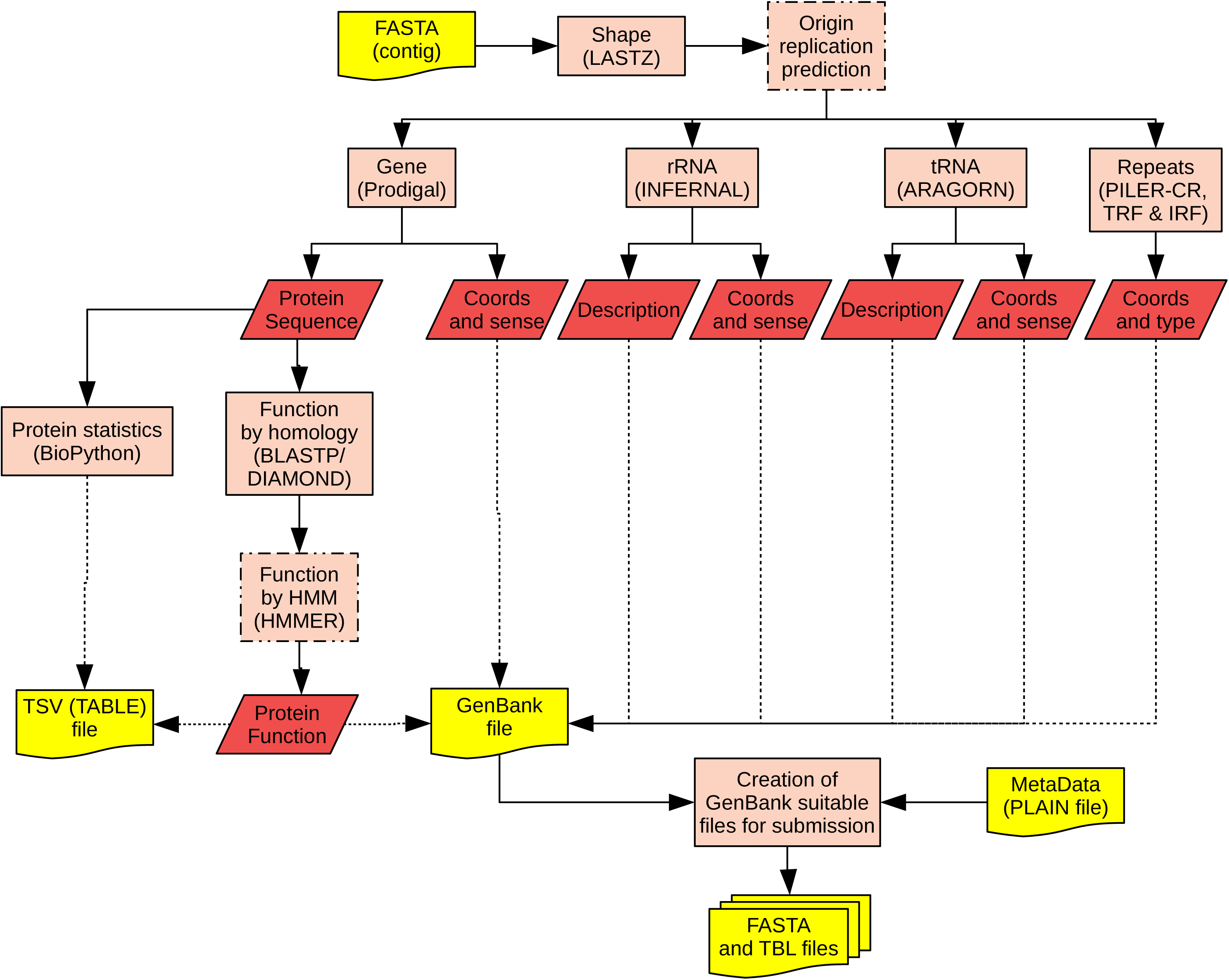
Flowchart of the VIGA pipeline. Orange rectangles represent the different steps of the program (among those, discontinuous-lined rectangles indicate optional steps; see main text). Red parallelograms indicate the relevant data that it is summarised in the output. Yellow rectangles with a wavy base stand for input and output files.

**Figure 5.**
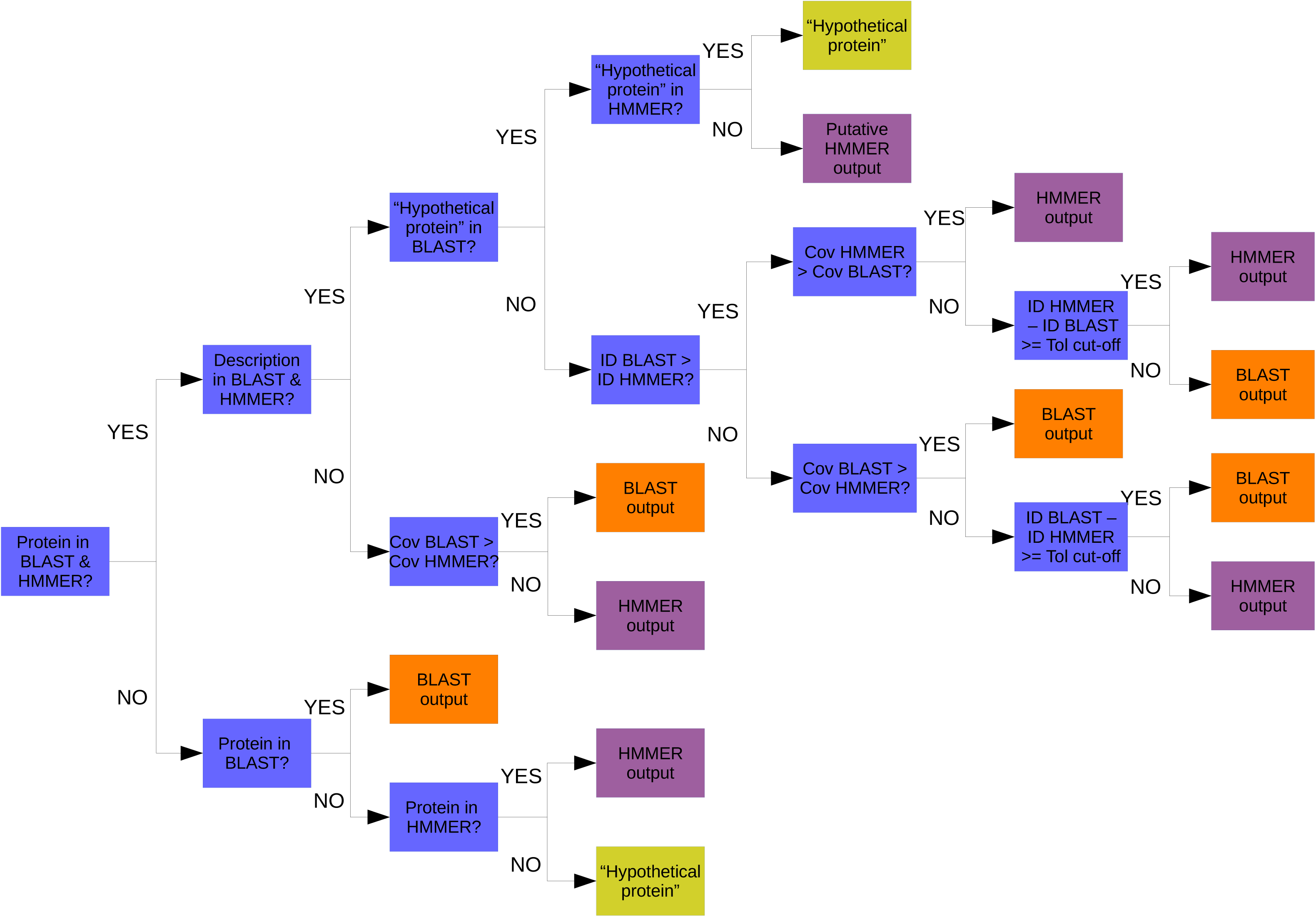
Flowchart of the decision tree algorithm. Blue rectangles represent steps in the decision tree. Orange and purple rectangles state optimal BLAST and HMMER solutions, respectively. Mustard coloured rectangles represent “hypothetical protein” decisions.

*Contig shape prediction.* VIGA requires a FASTA file containing a single or multiple sequences of viral contigs. Before running the gene prediction, VIGA launches LASTZ [55] to predict the circularity of every contig. In this case, a contig is defined as circular when the similarity between the initial and terminal fragment of the contig (by default the first and last 101 bp) is more than 95% and the length of such alignment covers more than 40%. When the contig is predicted as a circular, the software will predict the origin of replication based on iREP [56], which predicts the origin and terminus according to the cumulative GC skew.

*Gene prediction.* To predict genes in the contig, its length is checked and the most suitable program is run. If a contig is larger than 100,000 bp, Prodigal [57] is executed to predict the genes. If not, MetaProdigal [58] is launched to predict the genes. In both cases, when there are linear contigs, the programs are optimised to avoid predicting genes in regions near the closed ends of the contig. After the gene prediction, the coordinates and the protein sequence are saved.

*Function prediction.* Protein sequences are analysed using BLASTP [59] to predict its function according to homology. By default, BLASTP is run with default parameters (except for the *e*-value, which has been changed to 10^−5^ by default). However, an exhaustive BLASTP search could be performed using very strict values (a word size of 2, a gap open of 8, a gap extend of 2, the PAM70 matrix instead of BLOSUM62 and no compositional based statistics) to accurately identify proteins [60]. Alternatively, DIAMOND [47] can be used to predict protein function according to homology. For a more accurate protein function prediction, HMMER [48] can be executed to predict functions according to Hidden Markov Models with default parameters, except for the inclusion of an *e*-value cut-off of 0.001. To increase the protein function prediction speed, BLASTP can be launched using multiple threads and HMMER can run multiple jobs using GNU Parallel [61]. Both outputs are parsed independently according to identity, coverage, *e*-value and description to retrieve the protein function minimising the number of non-informative annotations as defined later.

*Decision tree algorithm.* If BLAST or DIAMOND were executed with HMMER to predict protein function, the BLAST/DIAMOND and HMMER outputs are processed using a decision tree to retrieve the description of every protein in the contig. For each protein, the existence of hits in both programs is checked. When the protein is detected in both BLAST and HMMER, non-informative annotations are detected searching for the expressions “hypothetical protein”, “uncharacterized protein”, “ORF”, “predicted protein”, “unnamed product protein” or “gp[Number]” in their BLAST and HMMER descriptions. If such a description is present in both proteins, the protein will be described as “hypothetical protein”. However if the “hypothetical protein” description is only present in BLAST, the consequent annotation retrieved by HMMER is considered as a valid one, and *vice versa*. In the scenario where the protein is not labelled as “hypothetical protein” in either BLAST or HMMER, it is checked if the percentage identity and coverage is higher in BLAST or in HMMER. Depending of these results, BLAST output or HMMER output is chosen accordingly (Fig. 5).

*rRNA prediction.* INFERNAL [49] is used altogether with the Rfam database [50] to predict the different ribosomal genes in every contig. In this case, INFERNAL hits are reported according to the gathering (GA) scores for every model.

*tRNA prediction.* ARAGORN [51] is launched to predict all tRNA and tmRNA sequences in every contig. After this step, the coordinates and the description of the tRNA are saved.

*CRISPR, tandem and inverted repeats prediction.* PILER-CR [52], Tandem Repeats Finder [53] and Inverted Repeats Finder [54] are used to detect CRISPR, direct tandem and inverted repeats in the contig, respectively.

*Output files.* After running all described steps, all saved information (contig shape, contig sequence, protein coordinates, protein sequences, protein descriptions, rRNA and tRNA coordinates, tRNA descriptions, and tandem and inverted repeats coordinates) is written to a GenBank file.

Additionally, the GenBank file is also converted to FASTA and TBL files after retrieving the metadata from a plain text file. The FASTA and the TBL files are suitable for GenBank submission. Optionally, a GFF file can also be created with this information.

### Benchmarking of VIGA

*Bioinformatic analysis.* 138 different viruses (52 bacteriophages, 72 eukaryotic and 10 archaeal viruses, and 4 virophages) which comprises 191 sequences (Additional file 1) were used to validate VIGA. Additionally, these sequences were also submitted to Prokka [6], RAST [5] and RASTtk [7] to compare their performance with VIGA. In this case, VIGA was launched in two different ways. First, VIGA was executed using BLAST [46] and HMMER [48] to predict protein function in the VIGA-Slow mode and then, launched using only DIAMOND [47] as the VIGA-Fast mode to predict protein function. In both cases, *nr* and UniProt databases were considered for DIAMOND/BLAST and HMMER, respectively.

*Statistical tests.* To evaluate the performance of VIGA, three different analyses were done. Firstly, to infer the accuracy and the precision of the number of viral coding sequences, general linear models were used. All linear models were forced to have intercept zero. The slope was used to measure the accuracy, while the R^2^ was used to measure the precision. Additionally, ANCOVA was used to compare the linear models. Secondly, the prediction quality of the coordinates of the viral coding sequences was evaluated by the F_1_ score, the precision and sensitivity, defined as

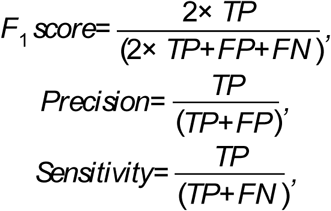

where TP indicates the number of true positives, FP the number of false positives and FN the number of false negatives. FDR and FNR were considered to measure the type I (i.e. false coordinates were considered as true coordinates) and the type II (i.e. true coordinates were considered as false coordinates) errors, respectively. To evaluate differences in the power of prediction of all programs, Kruskal-Wallis test was performed. In case that there were differences between programs, *post-hoc* tests using Nemenyi tests were performed. All statistical tests were carried out at an alpha level of 0.05 and were performed in R v. 3.4.1 [62] using the *HH* [63] and the *PMCMR* [64] packages.

### Case study: healthy human gut phageome

*Bioinformatic analysis.* VIGA was also tested on a metagenomic dataset using published data from the health human gut phageome [18]. This data set was downloaded from the SRA webpage (SRR codes: SRR4295172 – SRR4295175) and processed to retrieve contigs per sample. First, adapters were removed using Cutadapt 1.9.1 [65] and low-quality bases (lower than a PHRED score of 20 for a 4 bp sliding window) were trimmed using Trimmomatic [66]. All reads shorter than 30 bp were not considered for further analyses. All potential human reads were removed after being identified with Kraken v. 0.10.5 [67]. Contigs were assembled using metaSPAdes v. 3.10.0 [68] as recently the use of metaSPAdes was highly recommended to assemble metaviromes [69]. Assemblies of each sample were made non-redundant by an all-vs-all BLASTN [46] considering an *e*-value of 10^−6^. A contig was deemed redundant when it is shared 90% of its identity over 90% of the contig length. In these cases, the longer of the two contigs was retained. Non-redundant contigs over 1,000bp were processed using VirSorter [12] to generate a final data set of viral metagenome sequences. These contigs were annotated using VIGA in the two different ways described in the ‘Benchmarking of VIGA’ subsection and Prokka using 10 cores. Time benchmarking was performed using the *time* command in Linux only for VIGA and Prokka, as RAST and RASTtk are online genome annotation services.

*Statistical tests.* To evaluate differences in the power of prediction of all programs, Kruskal-Wallis test and *post-hoc* tests using Nemenyi tests were performed as described before. Moreover, to discard the effect of the length size of contigs as a potential factor of the power of prediction, Kruskal-Wallis tests were performed after classifying the contigs in three groups: “short” (<15 kb), “medium” (15 – 70 kb), and “long” (>70 kb). All statistical tests were carried out at an alpha level of 0.05 and were performed in R v. 3.4.1 [62] using the *HH* [63] and the *PMCMR* [64] packages.

## Declarations

### Ethics approval and consent to participate

Not applicable.

### Consent for publication

Not applicable.

### Availability of data and material

Source code of VIGA (and the wrapper for the Galaxy platform) is available for download at https://github.com/EGTortuero/viga, implemented in Python 2.7, and supported on Linux, under the GPL3 licence. The program is also available as at Docker image (https://hub.docker.com/r/vimalkvn/viga/).

### Authors’ contributions

LAD and SRS conceived the original idea. EGT and VV developed and wrote the VIGA software and the Galaxy wrapper. VV wrote the Docker integration of VIGA. EGT, LAD, SRS and CH designed the benchmark study. EGT and ANS tested VIGA against the validation benchmark. TDSS, EGT and ANS designed and run the case study. EGT, TDSS and SRS wrote the manuscript, with comments and editing by ANS, LAD, CH and RPR. All authors read and approved the final manuscript.

## Acknowledgements

EGT wants to thank Dr. Aditya Upadrasta for help in improving the software and Dr. Andrei Sorin Bolocan, Dr. Adam Clooney and Dr. Feargal J. Ryan for discussions.

## Additional files

**Additional file 1. List of the viruses used for the validation test (Excel file)**

